# The CCCH Zinc Finger Gene *PgCCCH50* from Pearl Millet Confers Drought Salt Tolerance through an ABA-Dependent PgAREB1-PgCCCH50 Module

**DOI:** 10.64898/2025.12.23.696222

**Authors:** Zheni Xie, Jie Zhu, Guohui Yu, Xixi Ma, Yi Zhou, Haidong Yan, Linkai Huang

## Abstract

Abiotic stresses like drought and salinity severely constrain crop productivity. Pearl millet (*Pennisetum glaucum* L.), a stress-resilient cereal of global importance, is ideal for deciphering genetic adaptation. Although CCCH zinc finger proteins play key roles in plant stress responses, members with multiple regulatory functions remain unknown in this crop. This study analyzed 53 *CCCH* genes in pearl millet, examining their phylogeny, structure, chromosomal localization, and promoter cis-elements. Expression profiling revealed seven *CCCH* genes responsive to multiple stresses. Functional characterization of *PgC3H50* showed that its overexpression in *Arabidopsis* conferred enhanced tolerance to drought and salt, evidenced by improved physiological parameters and upregulation of stress-responsive genes. We discovered that the ABA-responsive transcription factor PgAREB1(ABSCISIC ACID RESPONSIVE ELEMENT BINDING PROTEIN 1) directly binds to ABRE cis-elements in the *PgC3H50* promoter and activates its transcription, validated by yeast one-hybrid, dual-luciferase, and EMSA assays. This defines a novel PgAREB1-PgC3H50 regulatory module within the ABA-mediated stress signaling pathway. Our findings provide a valuable genetic resource for the pearl millet CCCH family and unveil a promising candidate gene and transcriptional regulatory module for engineering crops with improved tolerance to drought and salinity.

**Highlight:** This study reports the first genome-wide analysis of the CCCH family in pearl millet and identifies the novel PgAREB1-PgCCCH50 module as essential for salt and drought tolerance.

## Introduction

Abiotic stresses, such as drought and salinity, are critical constraints on plant growth and crop productivity (Ahmad et al., 2011). To cope with such adversities, plants have evolved intricate and interconnected regulatory networks that enable rapid and adaptive responses to environmental changes (Bavinck et al., 2014). Considerable efforts in genetic, biochemical, and molecular studies have identified numerous regulators involved in multilayer stress responses (Ait-El-Mokhtar et al., 2023).

Among these regulators, CCCH-type zinc finger proteins (ZFPs), characterized by tandem repeats of cysteine and histidine residues (typically C-X7/8-C-X5-C-X3-H) coordinated by zinc ions, have emerged as important players (Pi et al., 2021). CCCH proteins are widely implicated in plant stress defense, functioning as DNA-, RNA-, or protein-binding factors (Pomeranz et al., 2011). They have been identified across diverse plant species, with many showing responsive expression to abiotic stress. However, only a limited subset of these has been functionally validated. For instance, proteins such as AtSZF1/2, OsTZF1/5, PvC3H69/72, and SiC3H39 have been linked to stress tolerance in *Arabidopsis*, rice (*Oryza sativa* L), switchgrass (*Panicum virgatum* L.), and tomato (*Solanum lycopersicum* L.), respectively (Sun et al., 2007; Ilyas et al., 2022; Wang et al., 2024; Xie et al., 2019; Xu et al., 2023). Notably, while most studies focus on single-stress responses, growing evidence indicates that certain CCCH proteins, such as AtC3H15, GhZFP1, PeC3H74, HuTZF3, and IbC3H18, can confer tolerance to multiple concurrent stresses, including drought, salinity, and heat (Guo et al., 2009; Zhang et al., 2019; Lan et al., 2023; Xu et al., 2023; Andrés-Bordería et al., 2024). Despite these findings, the molecular mechanisms by which CCCH proteins orchestrate multi-stress responses remain poorly understood.

Abscisic acid (ABA) is a central phytohormone governing plant adaptation to drought and salinity. Several CCCH proteins operate in ABA-mediated signaling pathways. Examples include PvC3H69 in ABA-induced senescence, GhC3H20 in enhancing salt tolerance via interactions with PP2C phosphatases, PeC3H74 in ABA-mediated drought tolerance, and AtC3H3 in the upregulation of ABA-dependent genes under salt stress (Lan et al., 2023; Seok et al., 2024; Xie et al., 2021; Zhang et al., 2023). Nevertheless, current research predominantly centers on single-gene functions, leaving the exploration of CCCH proteins as components within larger molecular modules of the ABA pathway largely unclear.

Pearl millet (*Pennisetum glaucum* (L.) R. Br.) is a notable C4 cereal crop renowned for its resilience in arid and semi-arid regions, where it sustains millions under harsh climatic and soil conditions, including drought, salinity, and low fertility (Serba et al., 2016; Satyavathi et al., 2021; Daduwal et al., 2024). Unraveling the genetic basis of its abiotic stress tolerance is therefore crucial for breeding more resilient crops. However, the CCCH gene family has not been systematically identified in pearl millet, and its role in the plant’s stress regulatory networks is entirely unknown. In this study, we identified CCCH proteins in pearl millet responsive to multiple stresses using bioinformatics and RT-PCR analyses. Among these, we functionally characterized PgC3H50 and demonstrated its positive role in conferring both salt and drought tolerance. Furthermore, we revealed that the promoter of *PgC3H50* is directly bound and transcriptionally activated by the ABA-responsive transcription factor PgAREB1. Collectively, our findings establish the *PgAREB1–PgC3H50* module as a novel molecular component within the ABA-mediated regulatory pathway governing drought and salt stress responses. This study provides valuable genetic resources and a regulatory module for potential use in enhancing abiotic stress tolerance in crops.

## Materials and Methods

### Plant materials and growth conditions

Pearl millet ‘Tifleaf 3’ was used for gene cloning and expression analysis. After five minutes of disinfection with 0.1% sodium hypochlorite, pearl millet seeds were cleaned three times using distilled water. For germination, the seeds were spread on round Petri dishes (9×2 cm) with filter paper with a 12 h light/12 h dark photoperiod, 200 µmol/m^2^/s white light, and 60% relative humidity in a growth chamber at 28°C. Seven-day-old seedlings were transferred to half of Hoagland’s nutrient solution (pH 5.8). Afterwards, fourteen-day-old seedlings were subjected to heat treatment (45°C/40°C), cold stress (4°C), drought (20% PEG6000), salt (250 mM NaCl), and ABA (100 µM).

For *Arabidopsis* transformation, wild type (Columbia-0) was transformed via the floral dip method using Zhang’s Agrobacterium-mediated transformation method (Zhang et al., 2006). Forty-day-old seedlings of *Arabidopsis* wild type and transgenic lines were treated with drought and salt stress (300 mM NaCl) for phenotype and physiology analysis.

For the seed germination assay, the seeds of *Arabidopsis* wild type and *PgC3H50*-OE lines were disinfected with 50% (v/v) sodium hypochlorite and then sown on 1/2 MS and 1/2 MS with 150 mM NaCl or 300 mM Mannitol.

### Identification of CCCH proteins in pearl millet

The genome and gene annotation files of pearl millet ‘Tifleaf 3’, *Arabidopsis*, and rice were downloaded from the MilletDB (Sun et al., 2023) (http://milletdb.novogene.com/home/) and NCBI (https://www.ncbi.nlm.nih.gov/) databases, respectively. The Hidden Markov Model (HMM) of the CCCH conserved domain (PF00642) from the Pfam website (https://pfam.xfam.org/) (Mistry et al., 2021) and BLAST of homologous genes were used for candidate CCCH protein searching using the HMMER3.0 software (http://www.ebi.ac.uk/Tools/hmmer/search/hmmscan) (Finn et al., 2011). Conserved domains of candidate *CCCH* genes were confirmed using SMART (http://smart.embl-heidelberg.de/), HMMER, and NCBI Conserved Domains Search (https://www.ncbi.nlm.nih.gov/Structure/cdd/wrpsb.cgi) (Yang et al, 2020).

The physicochemical attributes of the candidate PgC3H proteins were predicted using Protparam (http://web.expasy.org/protparam/)(Duvaud et al., 2021). Moreover, the possible localization of PgC3H proteins within the cell was predicted using the BUSCA website (http://busca.biocomp.unibo.it/) (Savojardo et al., 2018).

### Construction of phylogenetic tree

The MEGA7.0 software ClustalW tool was used to align the amino acid sequences of CCCH proteins from pearl millet, *Arabidopsis*, rice (*Oryza sativa* L.), maize (*Zea mays* L.), and switchgrass (*Panicum virgatum* L.). The sequences were obtained from the NCBI database and Milletdb (Tamura et al., 2013). The phylogenetic tree was built using the neighbor-joining (NJ) method with 1000 bootstrap assessments. Lastly, the iTOL online tool (https://itol.embl.de/) was used to visually improve the NJ-phylogenetic tree (Letunic and Bork, 2021).

### Conservation motif and promoter analysis

Candidate CCCH conserved motifs were found utilizing the MEME online program (https://meme-suite.org/meme/tools/meme) with a limit of 10 motifs and a range of motif widths from 6 to 200, while keeping the remaining parameters at default values. The promoter sequences (approximately 2000 bp) of *CCCH* genes in pearl millet were extracted using TBtools software (https://github.com/CJ-Chen/TBtools) (Chen et al., 2020). Plant CARE (http://bioinformatics.psb.ugent.be/webtools/plantcare/html/) was used to identify possible cis-elements in the promoter region (Lescot et al., 2020).

### Gene cloning and vector construction

The CDS of *PgC3H50* was first cloned from the genome of pearl millet and then cloned into pENTR/D (Invitrogen, Carlsbad, CA, USA). The resulting construct was subsequently subcloned into pGreenII0800-LUC and pEarlygate103 using the LR reaction (NEB, Carlsbad, CA, USA). The primers used for vector construction and cloning are listed in Additional File S1.

### Subcellular location and transient transformation in Nicotiana benthamiana

To determine the subcellular location, the pEarleyGate103-*PgC3H50* vector was created by subcloning the CDS of *PgC3H50* into pEarleyGate103. pEarleyGate103-*PgC3H50* was then transformed into *GV3101* (Agrobacterium strain). A syringe was used to sporadically inject Agrobacterium carrying pEarleyGate103-*PgC3H50* and pEarleyGate103-*mGFP* into the mature leaves of 4-week-old *N. benthamiana*. A confocal laser scanning microscope (Zeiss LSM780 Exciter, Zeiss, Germany) was used to observe the GFP signal at 488 nm/bandpass of 505-525 nm.

Transient transformation of pEarlygate103-*PgAREB1* and empty vector (EV, pEarlygate103 only) in *Nicotiana benthamiana* was used for the functional analysis of *PgAREB1*. Leaves with transient overexpression of *PgAREB1* and EV were treated with 300 mM NaCl and 20% (w/v) PEG6000, respectively.

### Yeast one hybrid assay

The CDS of *PgABI5* and *PgAREB1* were subcloned into pGADT7, respectively. The promoter of *PgC3H50* was subcloned into pHIS2.1. Subsequently, the new pGADT7 vectors (pGADT7-*PgABI5*, pGADT7-*PgAREB1*) and pHIS2.1 vector (pHIS2.1-*pPgC3H50*) were introduced into the yeast strain Y187. Yeast were then cultivated on selection plates containing SD/-Trp-Leu-His and 120 mM 3-AT.

### Transcriptional activation assay

pGreenII 0800-*pPgC3H50* (the *PgC3H50* promoter) was generated for a transcriptional activation assay. The created vector and a control vector (pGreenII 0800-LUC vector only) were transformed into *GV3101* and then transiently transformed into leaves of *N. benthamiana.* Protein extraction was performed, followed by enzyme activity analysis, as stated by Yoo et al. (2007).

### RT-qPCR analysis

Leaves were sampled from pearl millet or *Arabidopsis* plants under stress treatment for RT-qPCR. All samples were immediately frozen in liquid nitrogen and ground into a powder. RNA was extracted using the Plant RNA Kit R6827 (OMEGA, USA), and cDNA was synthesized using the RT EasyTM II With gDNase kit (Chengdu Fuji Biotechnology Co., Ltd., Chengdu, China). The RT-qPCR assay was performed using the Real Time PCR easyTM-SYBR Green I kit (Chengdu Fuji Biotechnology Co., Ltd., Chengdu, China) on a Bio-Rad CFX Connect instrument, with the following calculated using the 2^−ΔΔCT^ method. The list of RT-qPCR primers is provided in Additional file S1.

### EMSA assay

*PgAREB1* was cloned into the pMAL-c5x vector for protein expression in *E. coli.* EMSA was performed as described by Xie et al. (2022).

## Results

### Identification and characterization of CCCH proteins in pearl millet

*CCCH* genes from pearl millet were searched using the Hidden Markov Model (Pfam: PF00642) released by the Pfam database. In pearl millet, 53 *CCCH* genes were found and designated as *PgC3H1* to *PgC3H53,* depending on their chromosomal locations. All 53 pearl millet CCCH proteins (PgC3Hs) were divided into 10 evolutionary branches numbered I to X (Figure. 1). The number of amino acids in the PgC3Hs ranged from 148 to 1776, with molecular weights ranging from 16238.06 Da to 192772.27 Da. The estimated isoelectric point (pI) ranged from 4.32 to 9.71. Subcellular localization predictions indicated that 45 PgC3Hs were found in the nucleus, accounting for 84.91% of the total. The remaining (15.09%) were distributed in other organelles, such as chloroplasts, membranes, and the outer membrane of chloroplasts (Additional file S2).

**Fig. 1.**
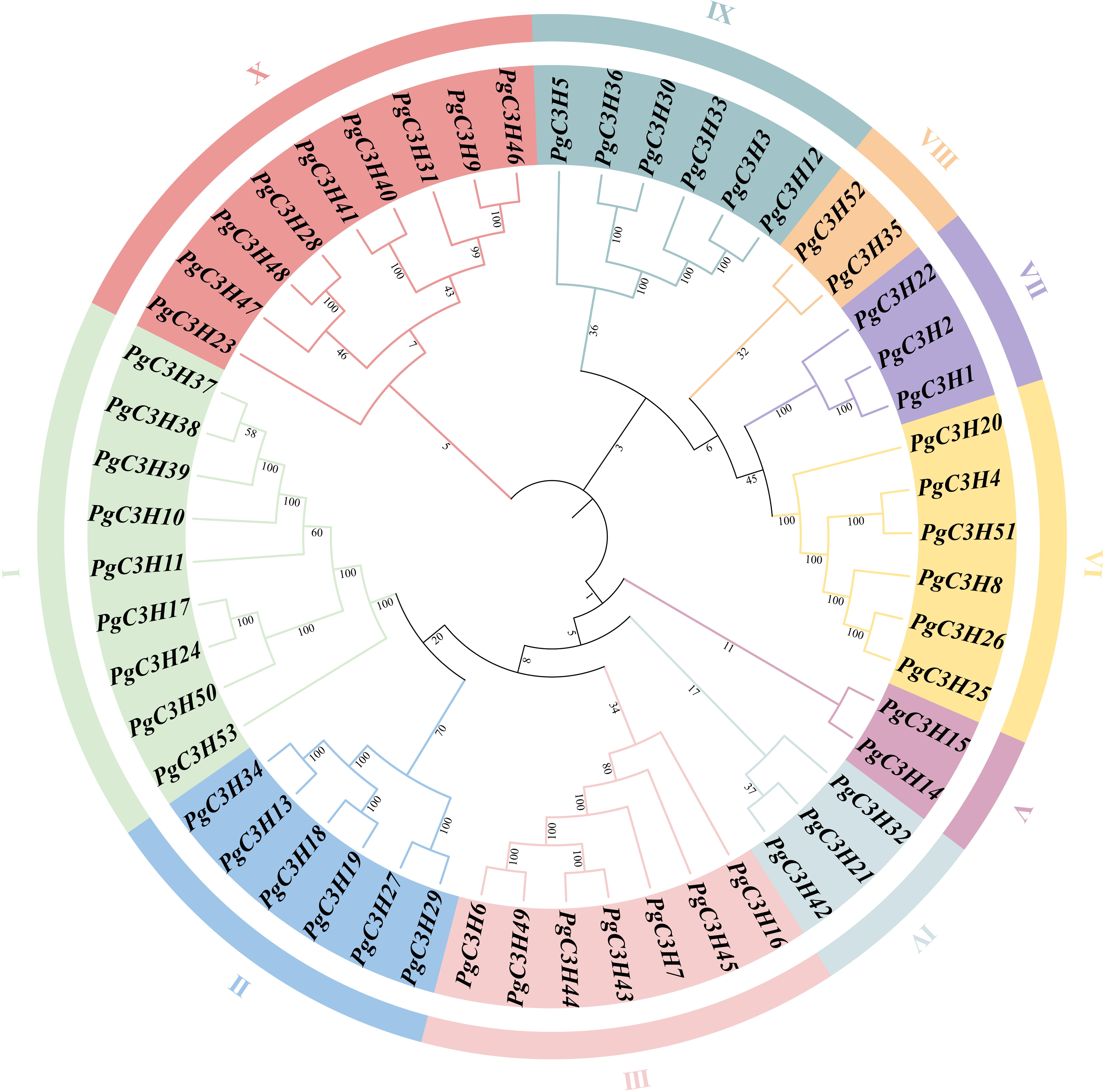
Phylogenetic tree analysis of *CCCH* genes in *Pennisetum glaucum.* The N-J method was used for constructing the evolutionary history in TBtools.

To examine the structural variation of *CCCH* genes in pearl millet (*PgC3Hs*), functional domains and exons-introns were analyzed in this study. Conserved motifs and genetic architectures showed similar patterns within the subclasses, consistent with the phylogenetic analysis. PgC3Hs belonging to the same evolutionary branch exhibited similar domain compositions and distributions, suggesting similar functions. Seven conserved domains were identified: Zf-CCCH, ANK, KH, zf-U1, WD-40, SAP, and RRM. Among the 53 PgC3Hs, eight had an RRM domain, four had an ANK domain, four had a KH domain, one had a WD-40 domain, and one had an SAP domain, with one to seven CCCH zinc fingers (Supplementary Fig. S1 A-B). For exon-intron analysis, as shown in Supplementary Fig. S2C, 11 (∼20.7%) CCCHs lacked exons, 10 CCCHs had no introns, and most Clade I members had 0-2 introns. Except for PgC3H20, Clade VI members had fewer introns, whereas other subfamilies contained more. Intron variations may occur through insertions or deletions, helping genes to develop new roles.

### Chromosomal location and gene duplication of CCCH family genes in pearl millet

The *PgC3Hs* were mapped to the pearl millet genome to better understand their chromosomal distribution. Forty-nine genes were spread over seven chromosomes (Supplementary figure S2), and four additional genes were found on the scaffolds. Among them, chromosome 6 contained the largest number of *PgC3Hs* (13), whereas chromosome 7 harbored only four. Chromosomes 1 and 5 contained seven *PgC3H* members, and three chromosomes (chr2, chr3, and chr4) contained six *PgC3Hs*.

Duplications were determined to better understand the expansion and evolution of *PgC3Hs*. Three pairs of *PgC3H*s showed tandem duplicates (TADs), and four pairs of segmental duplication genes (SEGs) were identified. All tandem and fragment repeat gene pairs had Ka/Ks ratios below 1, implying that *PgC3Hs* were in an essentially purified state, preventing the occurrence of harmful mutations during evolution and preserving gene function (Supplementary figure S2).

### Promoter and expression level analyses highlight 7 Clade-I CCCH genes associated with multiple abiotic stresses

To further understand the evolutionary connections between pearl millet and other plant species’ CCCH proteins, an evolutionary tree was created using CCCH proteins from five plant species: pearl millet, switchgrass, maize, rice, and *Arabidopsis* (Supplementary figure S3). The *PgC3Hs* in Clade I were homologous to *CCCH* genes associated with biotic or abiotic stress in *Arabidopsis*, maize, rice, and switchgrass (Figure 2). Cis-elements in promoters (∼2.0 kb) of *PgC3Hs* in Clade I were further investigated to determine their potential regulatory functions. As shown in Figure 3, nine types of cis-regulatory elements were searched in the promoter region of *PgC3Hs* These elements include the ABA Responsive Element (ABRE), Auxin Responsive Element, Low Temperature Responsive Element (LTRE), MeJA Responsive Element, SA Responsive Element, Drought-Inducible Element, GA-Responsive Element, Defense and Stress, and the Growth and Development Element. There were eight *PgC3Hs* in Clade I with multiple ABRE elements and eight *PgC3Hs* with MeJA elements, as well as four PgC3Hs that had defense and stress elements in their promoter regions (∼2.0 kb). Based on promoter analysis, it appears that Clade-I genes can respond to abiotic stress.

**Fig. 2.**
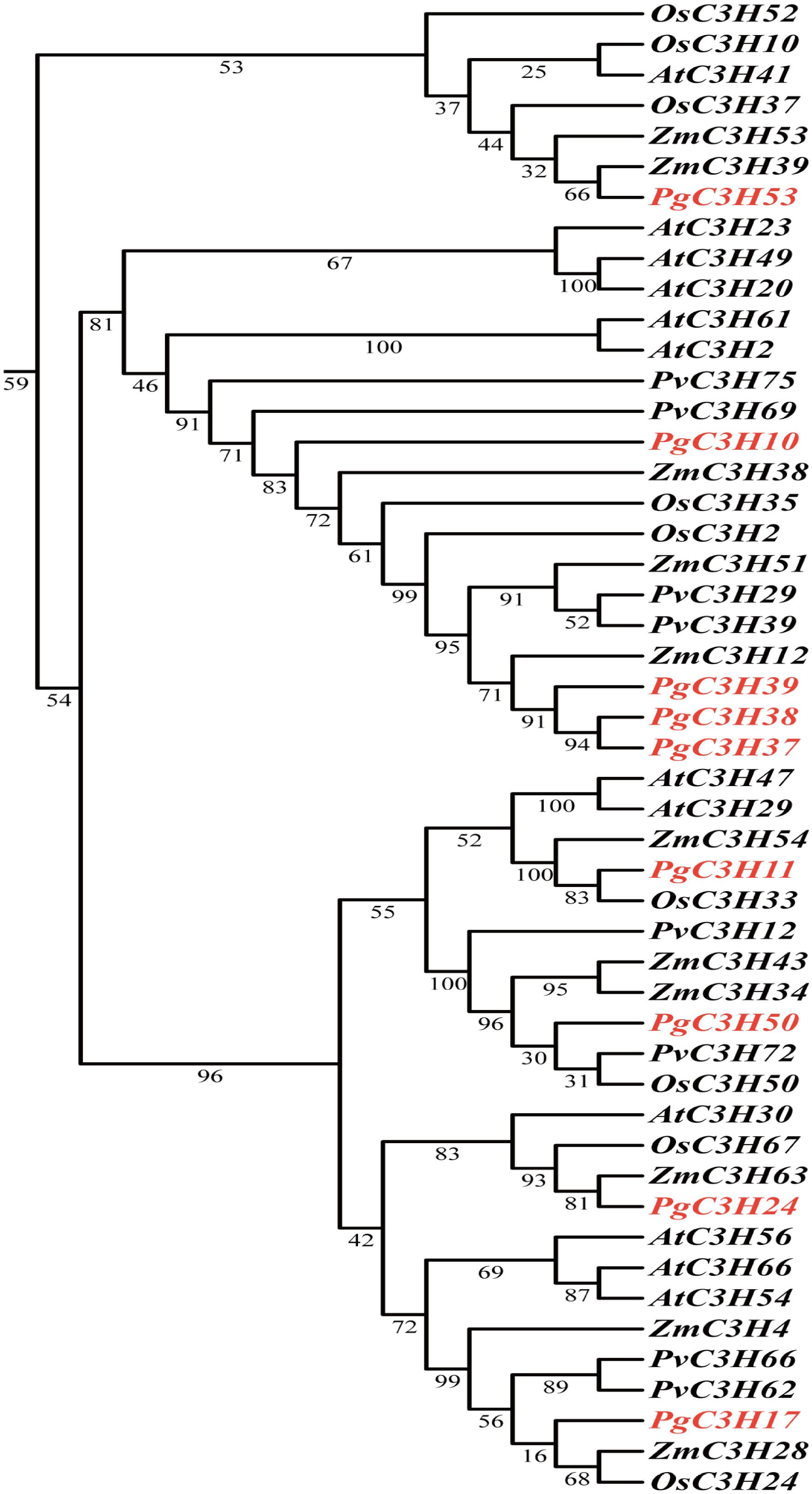
Phylogenetic tree of *CCCHs* in clade I in pearl millet and other plant species.

**Fig. 3.**
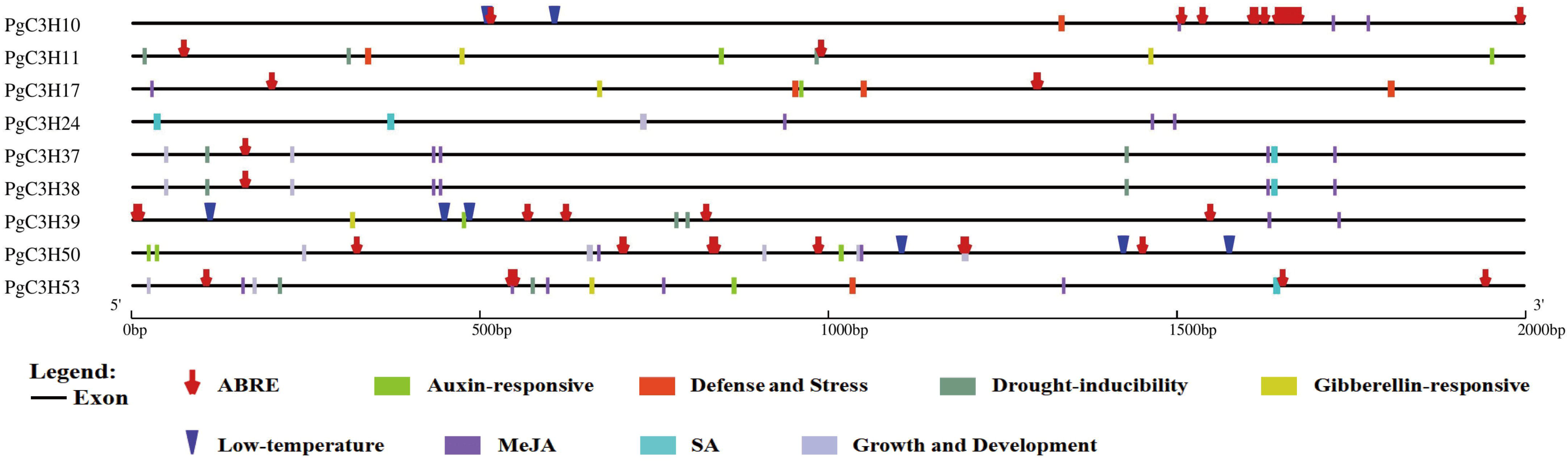
Cis-element analysis of the promoters of *CCCHs* in pearl millet.

Due to the complete sequence alignment rate (100%) between *PgC3H37* and *PgC3H38*, *PgC3H37* and the other seven *PgC3Hs* were further analyzed using RT-qPCR. As shown in Figure 4, multiple stress treatments significantly induced seven *PgC3Hs*, with the sole exception of *PgC3H39*, which only responded to PEG. *PgC3H10*, *PgC3H17*, *PgC3H24*, and *PgC3H53* exhibited notable responses to the four types of abiotic stress, except cold stress. Furthermore, *PgC3H11*, *PgC3H37*, and *PgC3H50* were dramatically induced by the three abiotic stresses. Specifically, *PgC3H11* expression was upregulated by heat, cold, and NaCl treatment. *PgC3H37* was induced by heat, PEG, and ABA, whereas the expression of *PgC3H50* dramatically increased under PEG, NaCl, and ABA conditions.

**Fig. 4.**
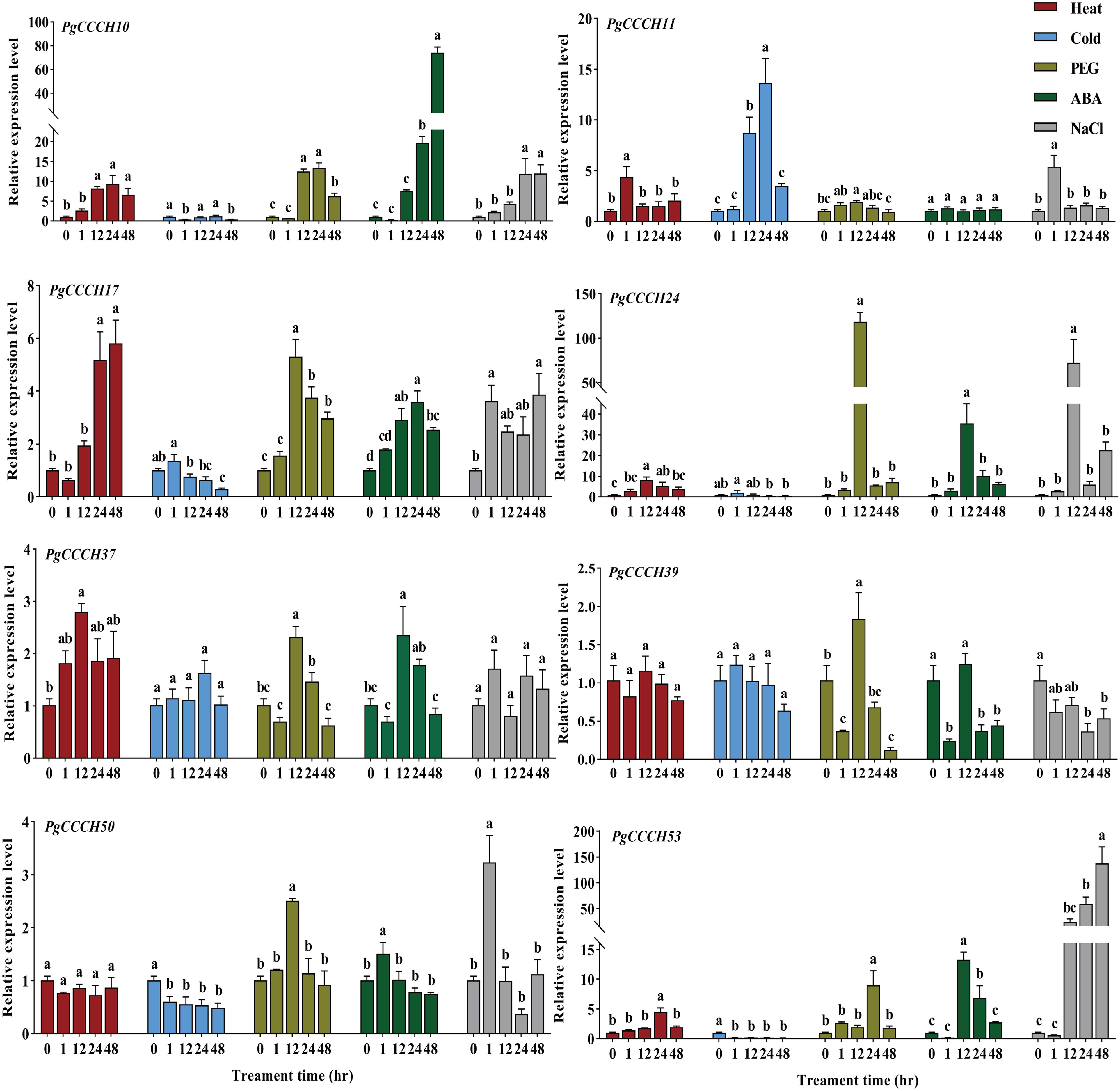
Relative expression levels of *CCCH* genes in pearl millet under heat (45D), cold (4D), 20% PEG6000 (w/v), ABA (100 µM), and NaCl (250 mM) treatments. Significant variations between treatments based on LSD levels are indicated by different letters. (P ≤ 0.05).

### PgC3H50 confers drought and salt tolerance in Arabidopsis

Then, *PgC3H50* was selected for further functional analysis. *PgC3H50 Arabidopsis* overexpression lines were generated to determine the function of PgC3H50 in regulating water stress. Three *PgC3H50*-overexpression (OE50) lines (OE50-2, OE50-3, and OE50-6) confirmed by RT-qPCR were exposed to drought stress. After a period of dryness lasting 12 d, the leaves of the wild type discolored green and became severely wilted, with even the tips of the flowers withering. In contrast, the leaves of all *PgC3H50*-overexpressing lines retained to 2-3 green leaves, and the flowers of all *PgC3H50*-overexpressing lines remained stretched. When re-watered for 6 days, wild-type plants failed to recover and almost died, whereas *PgC3H50* overexpression lines regained their vigor, producing more flowers and new green leaves (Fig. 5A-B). Also, the *PgC3H50*-overexpressing lines had much better survival rates than the wild type (Fig. 5C). Drought treatment resulted in significant changes in the physiological parameters. As shown in Figure 5D-E, compared to the WT, *PgC3H50*-overexpressing lines had significantly higher relative water content (RWC) but lower EL when subjected to 12 d of drought treatment followed by 6 d of watering recovery.

**Fig. 5.**
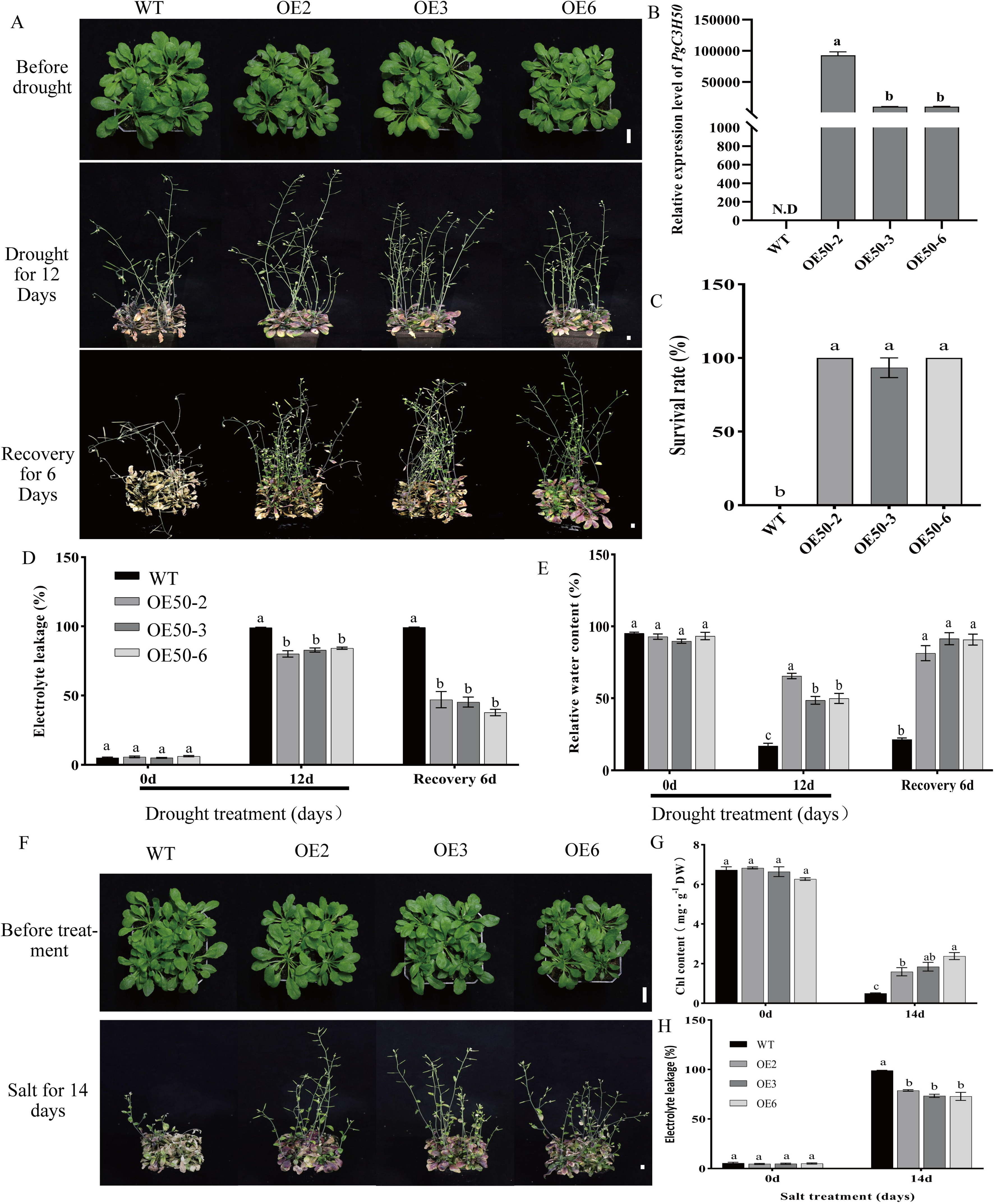
Characteristics of *PgC3H50* transgenic *Arabidopsis* under water deficit and salt stress. A. Phenotype of *PgC3H50-*overexpressing *Arabidopsis* and wild type under water deficit and recovery for 6 days. B. Relative expression of *PgC3H50* in *Arabidopsis* transgenic and wild-type lines. N.D. means not detected, and N.A C. Survival rates of *Arabidopsis* wild type and *PgC3H50-*overexpressing lines after 6 days of recovery. D. EL of *Arabidopsis* wild type and *PgC3H50-*overexpressing lines under water deficit and recovery for 6 days. E. Relative water content of *Arabidopsis* wild type and *PgC3H50*-overexpressing lines under water deficit and recovery for 6 days. F. Phenotype of *Arabidopsis* wild type and *PgC3H50-*overexpressing lines under salt stress. The white bars represent 1.5 cm length. G. Chl content of *Arabidopsis* wild type and *PgC3H50*-overexpressing lines with 300mM NaCl treatment. H. EL of *Arabidopsis* wild type and *PgC3H50-*overexpressing lines treated with 300 mM NaCl. The white bars represent 1.5 cm length. Significant variations between treatments based on LSD levels are indicated by different letters. (P ≤0.05).

To explore whether *PgC3H50* also contributes to salt tolerance, *PgC3H50*-overexpressing (OE50) lines (OE50-2, OE50-3, and OE50-6) and wild-type lines were grown under 300 mM NaCl. Under typical growing conditions, no visible phenotypic differences were observed between *PgC3H50*-overexpression plants and wild-type (WT) plants. However, after 14 days of treatment with 300 mM NaCl, the WT showed serious growth inhibition and almost died (Fig. 5F). Consistently, the Chl content of WT was significantly lower than that of *PgC3H50*-overexpression plants, whereas WT showed significantly higher EL than those in *PgC3H50*-overexpression lines (Fig. 5G-H).

Additionally, we evaluated the seed germination rates of both *PgC3H50*-overexpression lines and the wild type under salt or drought stress conditions. The results indicated that the germination rate of the *PgC3H50*-overexpression lines was significantly higher than that of the wild type under 150 mM NaCl or 300 mM mannitol treatment (Supplementary Figure. S4).

Furthermore, the relative levels of dehydration- and stress-responsive genes were determined using RT-qPCR. As shown in Figure 6, *PgC3H50* overexpression lines showed significantly higher expression levels of *DREB2A* and *DREB2B* than WT under salt or drought stress. Similarly, the expression levels of stress-responsive genes (*RD29A*, *RD29B*, and *RAB18*) in *PgC3H50* overexpression lines were significantly higher than those in WT when exposed to salt or drought, indicating that *PgC3H50* enhances plant stress tolerance (salt or drought) by controlling these key stress-responsive genes.

**Fig. 6.**
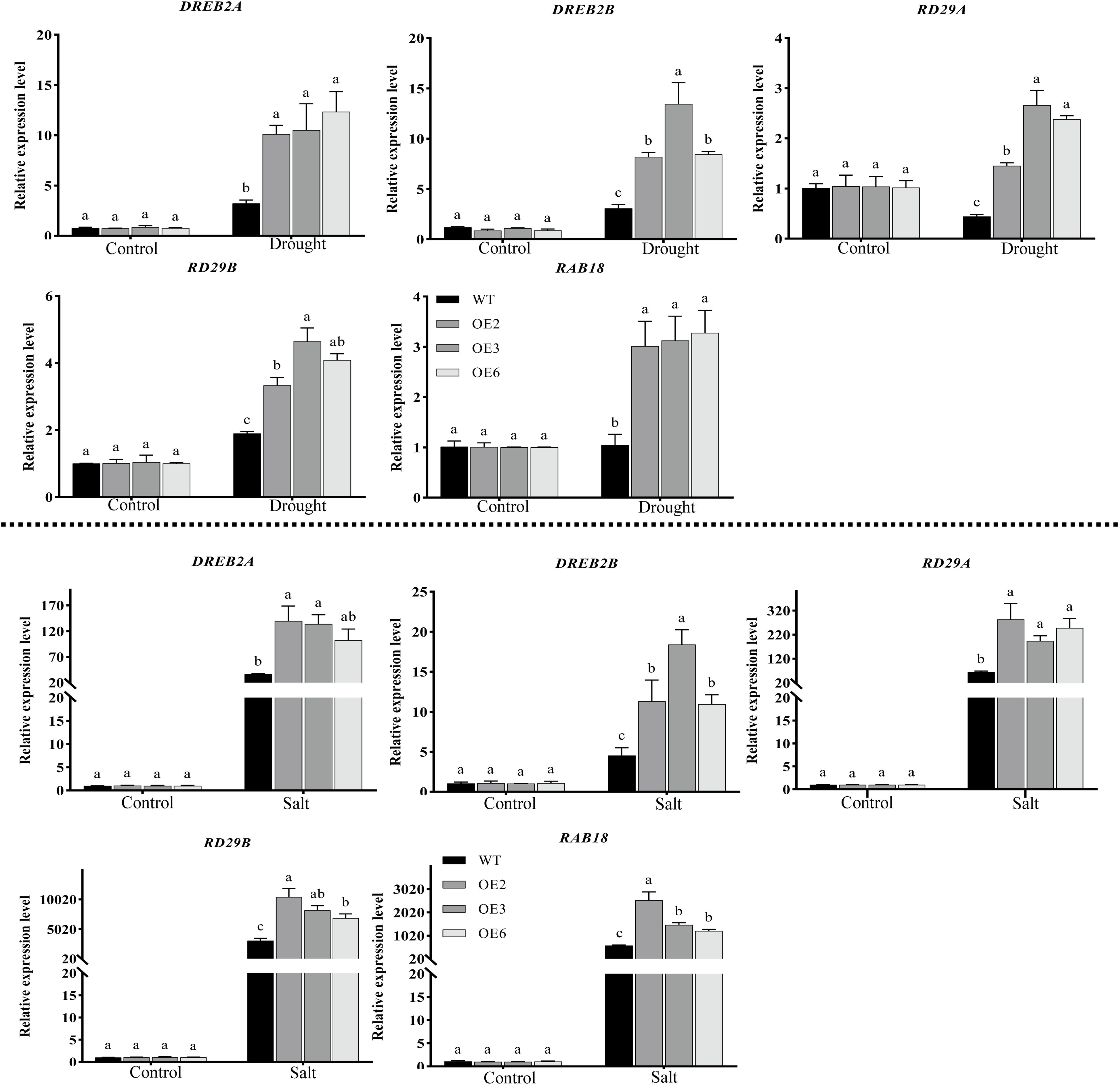
Stress response gene expression in *PgC3H50* transgenic lines and WT under salt or drought stress. Significant variations between treatments based on LSD levels are indicated by different letters. (P ≤ 0.05).

### Drought and salt stress regulation of PgC3H50 is associated with ABRE cis-element in its promoter

Subcellular localization was performed to determine the functional location of PgC3H50 in the cell. A GFP-fused PgC3H50 was constructed and transiently transformed into *Nicotiana benthamiana*. The aligned fluorescence of GFP indicated the cytoplasmic and membrane localization of PgC3H50, whereas the positive control (GFP only) was dispersed throughout the cell (Figure 7A). Considering that PgC3H50 may not be a typical transcription factor, we performed promoter analysis to further understand the common molecular mechanism of PgC3H50 in drought and salt stress regulation. Eight ABRE-responsive elements were identified in the promoter of *PgC3H50* (abbreviated as *pPgC3H50* hereafter) (Figure 7B). AREBs/ABFs (abscisic acid-responsive element binding proteins/ABRE-binding factors) are crucial for plant adaptation to several environmental stressors, particularly salinity and drought (Sarkar and Lahiri, 2013). Hence, we posit that the regulatory role of *PgC3H50* in response to salinity or drought is mainly involved in these ABRE-binding cis-elements. We further examined the promoter transcription activity of *PgC3H50* under salt and drought stress. Notably, after 3 h of salt treatment or 6 h of drought treatment, the transcription activity of the *PgC3H50* promoter was significantly higher than that before treatment; however, there was no difference in the transcription activity of the empty control, regardless of whether it was before or after stress treatment (Figure 7C-D). This result suggests that *PgC3H50* confers bi-stress defenses (drought and salt), which should be controlled by shared ABRE-binding proteins.

**Fig. 7.**
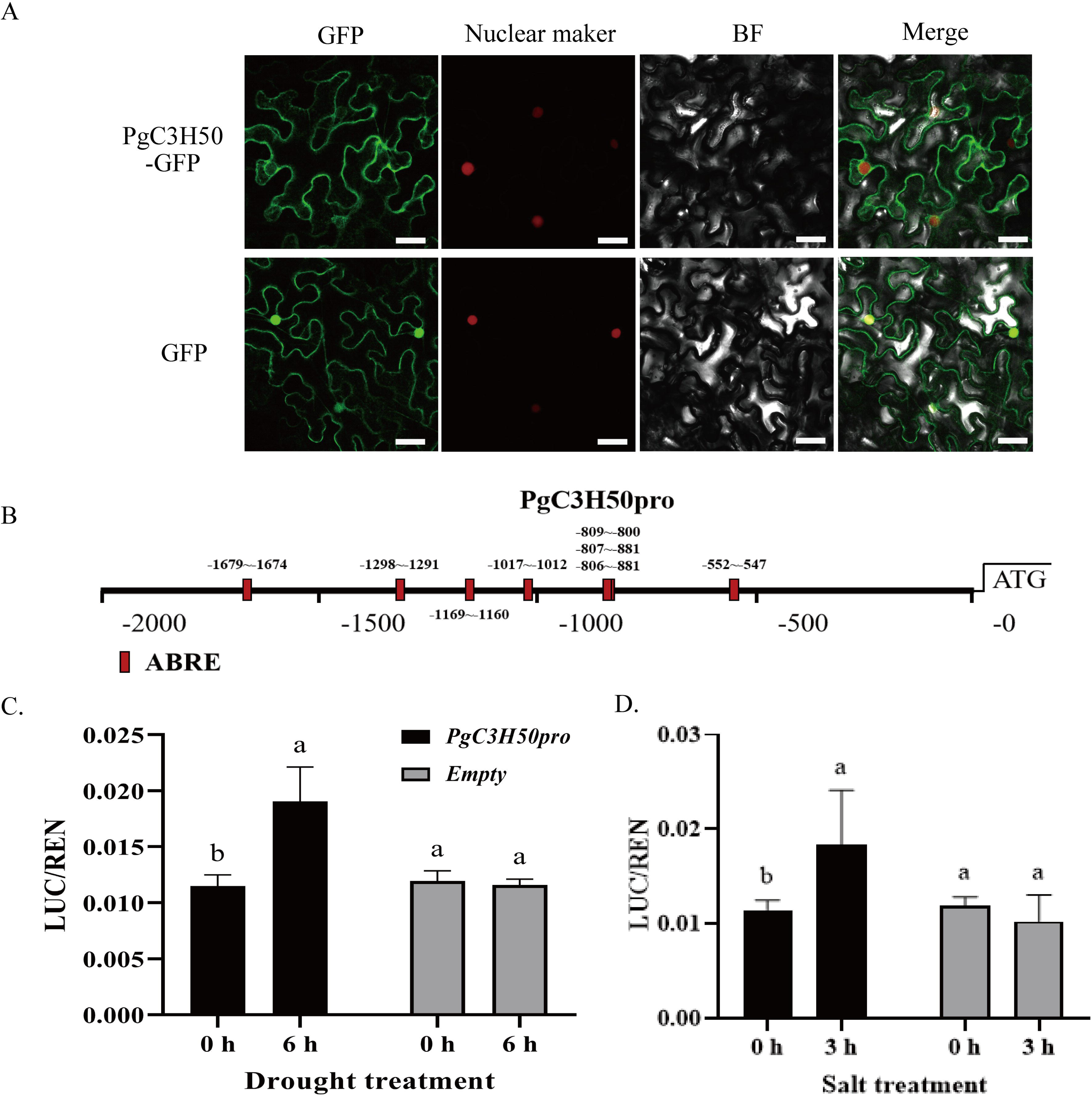
Subcellular localization of PgC3H50 and promoter transcription activity analysis of *PgC3H50* under drought and salt stress conditions. A. Subcellular location of PgC3H50. The green color indicates the localization of PgC3H50-GFP or GFP only, and the red color indicates the nuclear marker protein. BF, bright field; GFP, green fluorescent protein; white bars = 10 µm. B. Schematic diagram of promoter of PgC3H50; red boxes represent ABRE cis-elements. C-D. Promoter transcription activity analysis of PgC3H50 under drought and salt stress conditions. Significant variations between treatments based on LSD levels are indicated by different letters. (P ≤ 0.05).

### PgAREB1 directly binds to ABRE cis-element of pPgC3H50

ABI5 and AREB1 are well-known transcription factors that can directly bind to the ABRE cis-element of the target gene in response to drought and salt via the ABA-regulation pathway. The expression of *PgAREB1* and *PgABI5* in pearl millet under salt, drought, and ABA treatments was detected using RT-qPCR. Similar to *PgC3H50*, the expression of these genes was stimulated by drought, salt, and ABA (Figure 8). Notably, the expression of *PgABI5* was upregulated by approximately 15-and 30-fold under salt and drought stress, respectively.

**Fig. 8.**
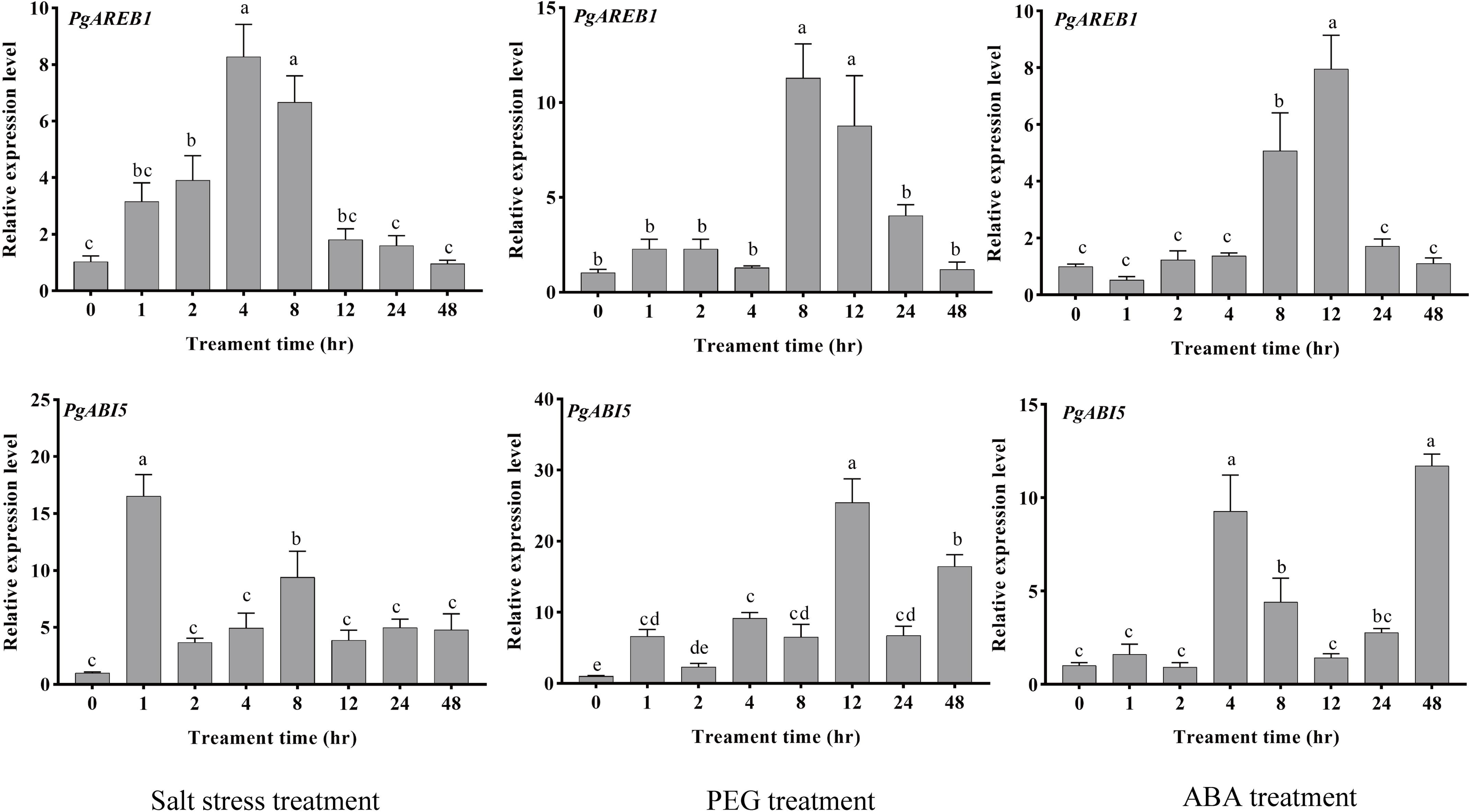
Relative expression levels of *PgAREB1* and *PgABI5* in pearl millet under salt (250 mM NaCl), 20% PEG6000 (w/v), and ABA (100 µM) treatment. Significant variations between treatments based on LSD levels are indicated by different letters (P ≤ 0.05).

To investigate whether *PgABI5* or *PgAREB1* interacted with *pPgC3H50*, a yeast one-hybrid assay (Y1H) was performed. As shown in Figure 9A, yeast cells co-transformed with PgABI5 and the promoter (∼2.0 kb) of *PgC3H50* or PgAREB1 and the promoter (∼2.0 kb) of *PgC3H50* survived better in SD/-TLH media with or without 3-AT supplements than the negative control. These results indicated that PgABI5 and PgAREB1 bind *pPgC3H50*.

**Fig. 9.**
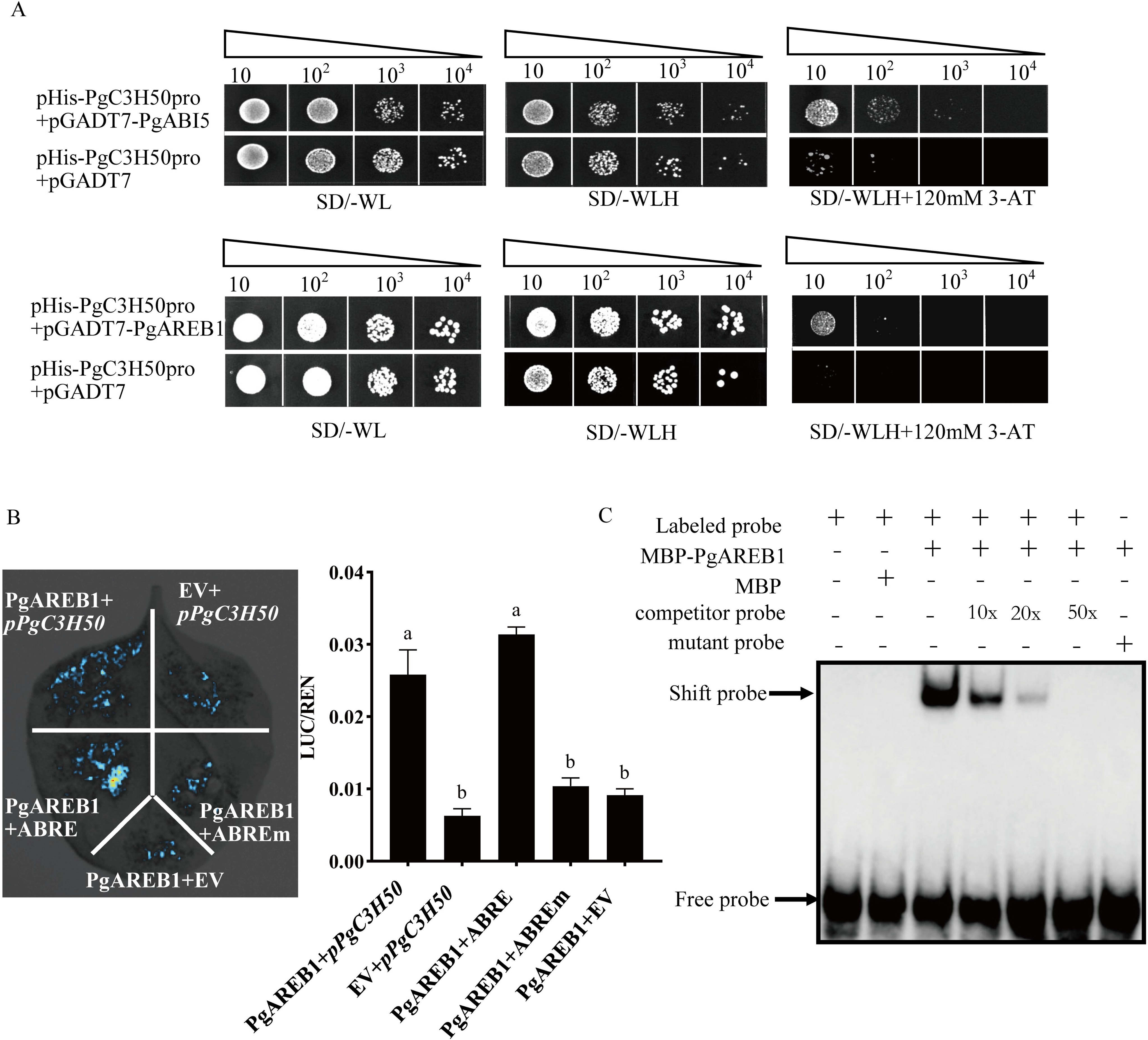
PgAREB1 directly binds to the ABRE cis-element of *pPgC3H50*. A. Yeast one hybrid of PgAREB1 and PgABI5 binding to the promoter of *PgC3H50*. SD denotes dropout amino acid selection medium; W, L, and H represent Trp, Leu, and His amino acids, respectively. 3-AT, Amitrol. B. Transcriptional activation assay in *planta.* PgAREB1 and LUC, driven by the PgC3H50 promoter, were co-expressed in *N. benthamiana.* The empty vector (EV) represented negative effector control. ABREm, a mutant of ABRE cis-element. Significant variations between treatments based on LSD levels are indicated by different letters. (P ≤ 0.05). All three experiments yielded similar results. C. EMSA assay for PgAREB1 binding to the promoter of *PgC3H50*.

Furthermore, a transcriptional activity assay was performed to verify whether the putative ABRE-binding factors could directly transactivate *PgC3H50*. As shown in Figure 9B, by co-expressing the effector (35S::PgAREB1) and the reporter gene (LUC) governed by *pPgC3H50* or ABRE-cis-element on *pPgC3H50*, it was shown that PgAREB1 trans-activated *PgC3H50*. However, when the ABRE-cis-element was mutated (3xACGTG to 3xATTTG) in *pPgC3H50*, PgAREB1 could not transactivate *PgC3H50*. To confirm direct binding, we performed an electrophoretic mobility shift assay (EMSA). The results showed direct binding of GST-PgAREB1 to the ABRE cis-element in the *PgC3H50* promoter, which was competitively reduced by increasing amounts of unlabeled probes and mutant probes (with CACGTG to AAAAAA mutation in the ABRE) (Figure 9C). Collectively, PgAREB1 directly transactivated PgC3H50 by binding the ABRE cis-element in *planta*.

### PgAREB1 enhances salt or drought tolerance

Transient overexpression of *PgAREB1* in *N. benthamiana* was performed to clarify its function under salt or drought stress, with empty vector transformation used as the control. In Supplementary Figure S5, under normal conditions, both *PgAREB1*-overexpressing and control leaves appeared emerald green and healthy, with no difference in EL. However, after 9 d of treatment with 300 mM NaCl or 20% PEG6000, the control leaves showed significant wilting and tissue damage, along with higher EL than *PgAREB1*-overexpressing leaves. These results indicate that *PgAREB1* plays a positive regulatory role in plant responses to drought and salt stress.

## Discussion

Pearl millet is a well-known millet species and the sixth most significant cereal crop globally, renowned for its ability to survive multiple abiotic stresses, such as salinity and drought (Varshney et al., 2017). Identifying genes that confer abiotic stress resilience in pearl millet is essential for breeding crops that can withstand stress.

CCCH zinc finger proteins (CCCH ZFPs) are key regulators of plant development and stress responses (Han et al., 2014). The proteins have been identified in over 10 plant species (Wang et al, 2008; Chai et al, 2012; Peng et al, 2012; Liu et al, 2014; Yuan et al, 2015; Chen et al, 2020; Hu and Zuo, 2021; Ai et al, 2022; Tang et al, 2023; Bao et al, 2024). However, CCCH proteins in pearl millet have not been identified or characterized. Here, 53 *CCCH* genes were identified in the pearl millet genome (Figure 1). The number is similar to Hordeum vulgare (Ai et al., 2022), but differs from other plant species. It appears that *CCCH* gene family composition varies by species.

CCCH proteins show functional diversity in abiotic stress regulation due to varying CCCH domains and extra domains like ANK, RRM, and KH. The ANK domain helps proteins stick together (Yuan et al., 2015; Islam et al., 2018), RRM aids mRNA binding (Vermel et al., 2002), and KH binds DNA/RNA (Lorković and Barta, 2002). AtTZF4, AtTZF5, and AtTZF6 interact with MARD1 and RD21A proteins (Bogamuwa and Jiang, 2016). PvC3H72 in *Panicum virgatum* acts as a transcriptional factor binding DNA for cold stress response (Xie et al., 2019). These findings show CCCH proteins can bind DNA or proteins, besides RNA (Pomeranz et al., 2010). We found several domains in pearl millet CCCH proteins, including SAP, ANK, RRM, WD-40, Zf-U1, and KH, suggesting functional diversity (Supplementary S1).

While many genes have been identified with multiple functional roles in stress regulation, reports on *CCCH* genes with dual or multi-stress regulation are limited. Cis-elements play a crucial role in the functional dissection and regulatory mechanisms of *CCCH* genes (Li et al., 2015). ABA-responsive elements (ABRE) are key cis-regulatory elements that regulate ABA-dependent gene expression. Abscisic acid (ABA) is crucial for controlling plant responses to external stressors, particularly during vegetative growth (Nakashima & Yamaguchi-Shinozaki, 2013). Drought-inducible elements are frequently located in the promoters of genes that are induced by drought stress, such as *RD22*, which is mediated by ABA (Abe et al., 1997). Similarly, the low-temperature-responsive element (LTRE), which contains an A/GCCGAC motif at its core, helps control the genes that respond to cold (Baker et al., 1994). Both ABRE and LTRE are important cis-elements that are responsible for abiotic stress responses. In our study, nine promoters of *CCCH* genes in Clade I contained multiple ABRE and LTRE cis-elements (Figure 3), and the increased expression of these genes during different abiotic stresses (Figure 4) suggested that many *CCCH* genes in pearl millet play key roles in response to multiple stressors. Notably, functional study indicated that *PgC3H50*, which has eight ABRE cis-elements in its promoter and is stimulated by salt, drought, and ABA, favorably regulates plant responses to salt and drought. (Figure 5). This was evidenced by a significantly elevated relative water content, reduced electrolyte leakage, and upregulated expression levels of stress-responsive genes in *PgC3H50*-overexpression plants subjected to drought or salt stress (Figure 6). Therefore, *CCCH* genes from pearl millet that respond to multiple stresses may function in different signaling pathways and serve as valuable candidates for the molecular breeding of stress-tolerant crops.

Additionally, several novel functional *CCCH* genes have been identified in species other than model plants, such as *Arabidopsis* and rice. For example, PvC3H72 in switchgrass has been identified as a novel cold-responsive transcription factor (Xie et al., 2019). The cotton novel gene GhZFP1 enhances salt and fungal disease resistance in transgenic tobacco (Guo et al., 2009). The tomato-specific SlC3H39 binds to the 3’-untranslated region of cold-responsive genes, leading to mRNA breakdown and post-transcriptional control of gene expression, thereby negatively modulating cold tolerance (Xu et al., 2023). These studies highlight the species-specific functions of *CCCH* genes. Interestingly, homologous *CCCH* genes can drive different functions in various species. In our study, PgC3H50 shared homology with *Arabidopsis* AtC3H47 (AtSZF1)/AtC3H29 (AtSZF2) and PvC3H72 in switchgrass (Supplementary figure S3&Figure 2). However, the homology between PgC3H50 and AtSZF1/AtSZF2 was only 45%. AtSZF1/AtSZF2 are known to regulate salt stress in *Arabidopsis*, but their involvement in drought regulation has not been further explored (Sun et al., 2007). In contrast, *PvC3H72* in switchgrass primarily responds to cold stress (Xie et al., 2019). Therefore, PgC3H50 in pearl millet exhibits distinct functions compared with AtSZF1/AtSZF2 or PvC3H72. The difference in the functional regulation of PgC3H50 could be due to the presence of multiple ABRE cis-elements (Figure 7), enabling it to participate in both salt and drought stress responses. Collectively, these findings highlight the probable target of PgC3H50 as a novel double-stress positive regulator in the cultivation of multi-stress crops.

ABRE-binding factors (AREB/ABF) are crucial for ABA signaling pathways and help plants adapt to various environmental stressors, especially salinity and drought (Fujita et al., 2011). AREB1 and ABI5, both members of the bZIP family, operate as ABRE-binding transcription factors to control downstream stress-responsive genes (Niyaz et al., 2017). The target genes of AREB1 have been revealed in many reports, including LEA class genes (e.g., *RD29B* and *RAB18*) (Niyaz et al., 2017) and other regulatory genes (e.g., *GBF3* and *RD20*) (Yoshida et al., 2010). However, no previous reports have confirmed that AREB1 or ABI5 binds to *CCCH* promoters, although several studies have demonstrated that CCCH proteins can regulate ABA-mediated pathways in plant stress or developmental processes (Xie et al., 2021; Zhang et al., 2023). In our study, *PgAREB1* and *PgABI5* exhibited expression patterns similar to *those of PgC3H50* under abiotic stresses (drought and salt), and PgAREB1 was found to directly bind and transactivate *PgC3H50* in *planta*, suggesting that PgAREB1 plays a critical role as an upstream regulator of PgC3H50 in the bifunctional stress regulation of both salt and drought (Figures 8, 9). Our study presents a novel “PgAREB1-PgC3H50” module essential for the resilience of pearl millet to salinity and drought stresses. Notably, the expression of *PgC3H50* was activated not only by dehydration and salt but also by ABA. Thus, whether “ PgAREB1-PgC3H50” can confer function in ABA stress should be further confirmed.

## Conclusion

This study presents the first genome-wide identification and characterization of the CCCH zinc finger protein family in pearl millet, revealing 53 members categorized into 10 subfamilies. Notably, seven genes from clade I were identified as responsive to multiple stressors, underscoring their potential role in complex environmental adaption. Specifically, PgC3H50 functions as a novel positive regulator conferring two tolerance to both drought and salt stress. Importantly, the novel candidate module “PgAREB1-PgC3H50” in which the transcription factor PgAREB1 directly binds to the ABRE cis-elements in the *PgC3H50* promoter in ABA-mediated drought and salt signaling pathways in pearl millet, as shown in Fig.10. These findings not only enhance the understanding of stress-responsive gene networks in pearl millet but also provide a promising genetic target (*PgC3H50*) and a precise regulatory mechanism for breeding crops with improved resilience to concurrent drought and salinity stresses.

**Fig. 10.**
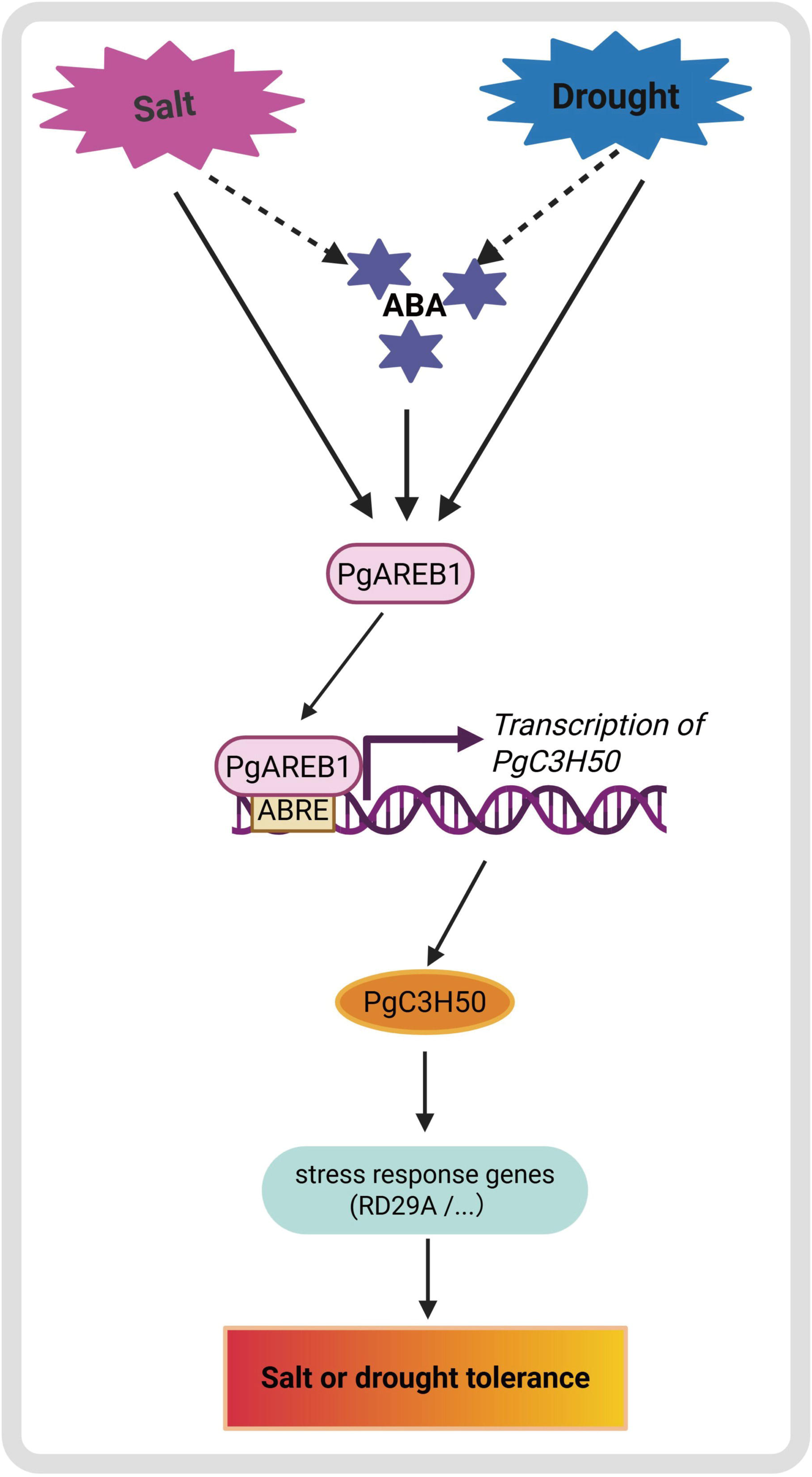
Transcriptional regulation of the “PgAREB1-PgC3H50” module in response to salt or drought stress.

## Acknowledgements

We thank Prof. Bin Xu from Nanjing Agricultural University for contributing several vectors to our research.

## Author contributions

Zheni Xie and Linkai Huang developed the research and experimental designs. Jie Zhu, Xixi Ma, Guohui Yu, Yi Zhou and Haidong Yan performed the experiments. Zheni Xie and Jie Zhu analyzed all the data. Zheni Xie wrote the manuscript. All the authors have read and approved the final manuscript.

---

Supplementary figure S1: Gene structures and functional motifs of *CCCHs* in pearl millet. A. Evolutionary relationship and functional motif distribution of *CCCHs*. B. Functional motifs of CCCHs. C. Gene structures of *CCCHs*.

Supplementary figure S2: Chromosome distribution and gene duplication of *CCCHs* in pearl millet. A. Synteny analysis of *CCCH*s in pearl millet. B. Gene number of TADs and SEGs. C. Ka/Ks analysis of TADs and SEGs.

Supplementary figure S3: Phylogenetic tree of 53 CCCHs in switchgrass, rice, maize, Arabidopsis, and pearl millet.

Supplementary figure S4: A. Seeds germination of *Arabidopsis* PgC3H50-overexpression lines and wild type under 150mM NaCl. B. Seeds germination rate of *Arabidopsis* PgC3H50-overexpression lines and wild type under 150mM NaCl. C. Seed germination of *Arabidopsis PgC3H50*-overexpression lines and wild type under 300mM mannitol. D. Seeds germination rate of *Arabidopsis PgC3H50*-overexpression lines and wild type under 300mM mannitol.

Supplementary figure S5:PgAREB1 enhances salt and drought tolerance. A. Phenotype of transient *PgAREB1*-overexpressing and *EV* tobacco leaves under salt or PEG treatment. B. Relative expression level of *PgAREB1* in transient *PgAREB1*-overexpressing tobacco leaves and *EV* tobacco leaves. C. Physiological parameters of transient *PgAREB1*-overexpressing and *EV* tobacco leaves under salt or PEG treatment. *EV*, empty vector with GFP only.

Supplementary Table S1 : Summary of the basic characteristics of *the CCCH* gene family in pearl millet.

Supplementary Table S2: Primers used in this study.

